# Revisiting *bicoid* function: complete inactivation reveals an additional fundamental role in *Drosophila* egg geometry specification

**DOI:** 10.1101/2023.12.08.570753

**Authors:** Stefan Baumgartner

## Abstract

**Introduction:** The *bicoid* (*bcd*) gene in *Drosophila* has served as a paradigm for a morphogen in textbooks for decades. Discovered in 1986 as a mutation affecting anterior development in the embryo, its expression pattern as a protein gradient later confirmed the prediction from transplantation experiments. These experiments suggested that the protein fulfills the criteria of a true morphogen, with the existence of a homeodomain crucial for activation of genes along the anterior-posterior axis, based on the concentration of the morphogen. The *bcd* gene undergoes alternative splicing, resulting in, among other isoforms, a small and often neglected isoform with low abundance, which lacks the homeodomain, termed *small bicoid* (*smbcd*). Most importantly, all known classical strong *bcd* alleles used in the past to determine *bcd* function apparently do not affect the function of this isoform.

**Results:** To overcome the uncertainty regarding which isoform regulates what, I removed the *bcd* locus entirely using CRISPR technology. *bcd^CRISPR^* eggs exhibited a short and round appearance. The phenotype could be ascribed to *smbcd* because all *bcd* alleles affecting the function of the major transcript, termed *large bicoid* (*lgbcd*) showed normally sized eggs. Several patterning genes for the embryo showed expression in the oocyte, and their expression patterns were altered in *bcd^CRISPR^* oocytes. In *bcd^CRISPR^* embryos, all downstream segmentation genes showed altered expression patterns, consistent with the expression patterns in “classical” alleles; however, due to the altered egg geometry resulting in fewer blastoderm nuclei, additional constraints came into play, further affecting their expression patterns.

**Conclusions:** This study unveils a novel and fundamental role of *bcd* in shaping the egg’s geometry. This discovery demands a comprehensive revision of our understanding of this important patterning gene and prompts a reevaluation of past experiments conducted under the assumption that *bcd* mutants were *bcd^null^*-mutants.

## Introduction

The *bicoid* (*bcd*) gene in *Drosophila* and its product, the morphogenetic gradient have fascinated researchers for decades (reviewed by (Baumgartner, 2018). Historically, the gene was identified in a screen for maternal mutants responsible for anterior development (Frohnhöfer and Nüsslein-Volhard, 1986). It was demonstrated that cytoplasmic transplantations from anterior cytoplasm could elicit a response at the recipient’s egg and generate anterior structures, indicating the presence of a substance capable of inducing anterior structures. In the same year, another group cloned the *bcd* gene using a DNA homology-based approach (Frigerio et al., 1986) confirming its anterior expression and the formation of an mRNA gradient at blastoderm stage. Two years later, the Bcd protein gradient was demonstrated through antibody staining (Driever and Nüsslein-Volhard, 1988). However, the previously demonstrated mRNA gradient (Frigerio et al., 1986) was neglected and a new model was presented, the SDD model (Driever and Nüsslein-Volhard, 1988), reviewed by (Baumgartner, 2018). In this model, S stands for synthesis, D diffusion and D for degradation, representing a simplified model for the diffusion of Bcd throughout the cytoplasm, based on the French-Flag model (Wolpert, 1969), itself rooted on Alan Turing’s reaction–diffusion model (Turing, 1952). Bcd became the first demonstrated morphogen to fulfill all the predictions of the properties of a diffusible factor synthesized at a source and eliciting different cell fates based on concentration. This served as a paradigm for morphogens in biological textbooks for over two decades. However, in 2007, doubts were raised about the diffusion constant being too low to reach posterior positions (Gregor et al., 2007b), a concern that was eventually corrected and found to be sufficiently high (Abu-Arish et al., 2010; Durrieu et al., 2018; Sigaut et al., 2014). Notably, the SDD model was deemed too simplistic. While initial support for broad posterior diffusion came from studies using fluorescent dextran particles injected at the anterior pole (Gregor et al., 2005), it took some time to refute this notion. Bcd movement was revealed to be limited to the cortex (Baumgartner, 2018; Cai et al., 2017; Gregor et al., 2007b), with the inner yolk acting as a barrier, preventing any movement. Only when the internal barrier, the yolk, was compromised using microtubule-degrading drugs did Bcd movement conform to the SDD model and the dextran particle movement, showing broad posterior movement in the inner yolk (Cai et al., 2017).

In 2009, a new model of gradient formation, the ARTS model (A standing for active, R for RNA, T for transport, S for synthesis) was proposed (Baumgartner, 2018; Fahmy et al., 2014; Lipshitz, 2009; Spirov et al., 2009). This model integrated an old observation from 1986, where an mRNA gradient was noted (Frigerio et al., 1986), suggesting that the *bcd* mRNA was transported from the anterior pole with the help of microtubules (MTs) to the posterior, forming a gradient. An essential step in gradient formation was reported to occur during mid/late nuclear cycle 14 when the mRNA was transported from the basal to the apical side, establishing a gradient extending to at least 40% of the egg length (Fahmy et al., 2014; Frigerio et al., 1986; Spirov et al., 2009). This mRNA gradient was proposed to serve as the source for translation of the *bcd* mRNA, forming the Bcd protein gradient.

Soon after, another report (Little et al., 2011) challenged this idea, asserting that the mRNA gradient was nearly absent and that 90% of the mRNA concentrated within the anteriormost 20% of the embryo. However, this report had a significant flaw as it relied on insufficiently fixed embryos, leading to extensive tissue loss. Notably, the preservation of the cortical cytoplasm, also termed periplasm, was compromised, as the embryos did not survive the steps of FISH very well, exhibiting abrasion of the periplasm up to the layer of the nuclei. This resulted in a misleading conclusion that the *bcd* mRNA was degraded uniformly at early nc14, with no basal-to-apical movement of the mRNA detected. As no periplasm was present, no long periplasmic mRNA gradient formation was observed. The failure to properly preserve the integrity of tissues raises further questions about the overall preservation of mRNA within the embryos discussed in this report.

All classical *bcd* mutants originate from an initial screen for maternal mutants, which yielded 11 different *bcd* alleles (Frohnhöfer, 1987; Frohnhöfer and Nüsslein-Volhard, 1986). Complementary alleles were obtained from screens conducted at later time points (Luschnig et al., 2004; Seeger, 1989; Seeger and Kaufman, 1990). Notably, all mutant alleles were derived from EMS screens, resulting in single base pair changes in most cases (Fig. 1D). However, due to the presence of the homeodomain (Fig. 1B) dominating the gene’s function, strong alleles exhibiting anterior patterning defects were favored, while the weaker alleles with milder to no anterior phenotype were often overlooked. Over time, many original *bcd* mutant stocks failed to yield homozygous *bcd* females, likely due to additional lethal hits. Consequently, *bcd*^-^ mutant mothers could only be obtained in *trans* to another *bcd* allele, hampering structure-function analyses. This limitation led a group creating a well-defined *bcd* mutant and developing a strong CRISPR-based *bcd* allele using a MiMIC cassette inserted into the first intron (Fig. 1D), with the idea that the splice event would create *trans*-splicing between exon 1 of *bcd* and the first available exon on the MiMIC cassette (Huang et al., 2017). This event resulted in a short and truncated Bcd protein without the homeodomain. The resulting *bcd* mutant faithfully recapitulated the appearance of the strongest *bcd* alleles (Huang et al., 2017). In a recent study (Fernandes et al., 2022), a complete removal of the *bcd* locus based on CRISPR technology was achieved. However, no in-depth analysis was conducted using this allele.

**Figure 1.**
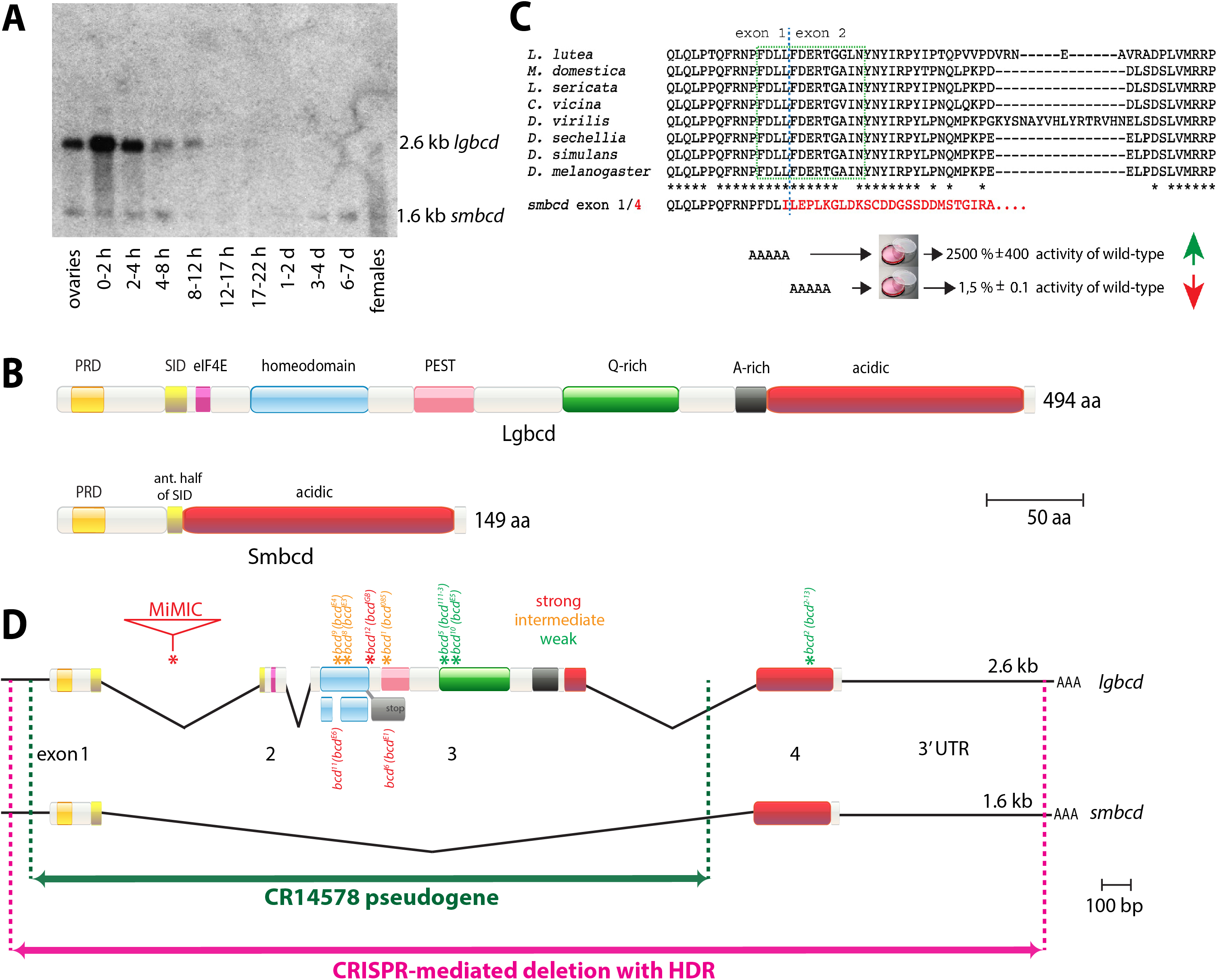
Genomics of the *bcd* locus. (A) Original *bcd* Northern filter from (Berleth et al., 1988), rescanned and adjusted to denote the 1.6 kb *smbcd* transcript that was not visible in the original publication due to printing constraints. *smbcd* is considerably less abundant than *lgbcd* but is expressed much broader during development. Stages and intervals where polyA^+^ RNAs were collected are indicated at the bottom. B) Top, domain structure of large Bicoid with annotated domains; Bottom, domain structure of small Bicoid. Notably, Smbcd lacks the DNA-binding activity of the homeodomain (bright blue) and the COOH-terminal part of the self-inhibitory domain (SID; yellow), compare in Fig. 1C. C) Sequence of the SID domain from different insects (from top: *Lonchoptera lutea, Musca domestica, Lucilia sericata, Calliphora vicina, Drosophila virilis, Drosophila sechellia, Drosophila simulans, Drosophila melanogaster*) aligned, with 100% identities annotated by asterisks below. Blue dashed line denotes borders of exon 1 and exon 2, green dashed box denotes core part of the SID. The amino acid sequence of Smbcd, where exon 1 and 4 (in red) are joined, is indicated below. AAAAA denotes mutagenized amino acids, and the resulting changes in Bcd activity are indicated in percentages compared to wild-type Bcd (Zhao et al., 2003). The fusion of exon 1 to exon 4 to create *smbcd* has been documented in *Drosophila melanogaster*, all Drosophilidae and *Lucilia sericata* so far. D) Molecular map of the *bcd* locus, illustrating genome organization, strengths, and mapped molecular lesions of *bcd* mutants, extent of the homology to *bcd* pseudogene CR14578, and outline of the CRISPR-mediated deletion with HDR. The *bcd* gene exhibits alternative splicing, generating five transcripts, two of which - *large bicoid* (*lgbcd*) and *small bicoid* (*smbcd*) - are displayed. In the latter, exons 2 and 3 are omitted. Three other seemingly minor alternative splice events were also reported (FlyBase). Mutants with mapped molecular lesions (Struhl et al., 1989) fall into three classes, classification according to (Frohnhöfer, 1987): strong (red) and intermediately strong (orange) alleles reveal lesions, primarily as point mutations (asterisks) around the homeodomain (light blue). A MiMIC allele results in a truncated protein due to integration into the first intron. Three alleles are weak ones (green) and show lesions outside the homeodomain. Allele names are indicated above the lesion as FlyBase name, followed by the original name from (Frohnhöfer, 1987). The 3L-region in the heterochromatin harbors a large stretch of *bcd* homologous sequences with almost 100% DNA sequence identity, annotated as an inactive pseudogene CR14578 (green). The extent of the CRISPR-mediated deletion of the genomic *bcd* region and subsequent replacement with an *eyeless*-DsRed cassette is shown in pink.

The first molecular description of *bcd* (Berleth et al., 1988) revealed that the gene produced two types of mRNA through alternative splicing. The large splice product was the focus of the study due to its large abundance and harboring the homeodomain, while the small transcript was shown to be generated through a splice event omitting exon 2 and 3 (and thus the homeodomain), joining exon 1 and 4. Among the many cDNAs isolated from the locus, one cDNA (c53.46.5) confirmed the proposed alternative splicing event. Unfortunately, due to the small transcript’s low abundance and constraints during the printing process, the band of the small transcript was no longer visible in the actual reproduction of the Northern blot. Since the small transcript appeared to lack function, being without the homeodomain, it was virtually neglected over the last three decades. To distinguish between the two isoforms, they are now referred to as *large bcd* (*lgbcd*) and *small bicoid* (*smbcd*).

Since the function of *smbcd* is likely unaffected in all classical *bcd* alleles, I aimed to create a clear situation to understand the comprehensive role of the entire *bcd* locus. To achieve this, the entire *bcd* locus was deleted using CRISPR technology. Complete removal of *bcd* uncovered a new function for *bcd* in shaping the egg geometry, resulting in eggs that are short but round with fewer blastoderm nuclei. This mechanism is likely to be controlled by *smbcd* and not by *lgbcd*, thereby adding a new and fundamental function for *bcd* during oogenesis.

## Results

In the past four decades, a plethora of data on *Drosophila bicoid* (*bcd*) function, appearance and its definition as the first morphogen have been generated. This data largely relied on the analysis of the isoform containing the homeodomain, as reviewed by (Baumgartner, 2018; Grimm et al., 2010; Huang and Saunders, 2020; Irizarry and Stathopoulos, 2021; Wieschaus, 2016).

While the original publication of the cloning and the subsequent molecular analysis uncovered precious details (Berleth et al., 1988), an essential detail went unnoticed due to print constraints during the reproduction of the original Northern analysis of the *bcd* gene. Specifically, a small and considerably less abundant isoform of the *bcd* transcript, a 1.6 kb transcript termed *small bicoid* (*smbcd*) was simply not visible in the figure. Moreover, this 1.6 kb transcript was barely mentioned in the text due to the significance of the larger 2.6 kb transcript, now termed *large bicoid* (*lgbcd*). As a consequence, *smbcd* was largely forgotten for the next 35 years, with the exception of a 2013 publication by (Rodel et al., 2013) where it showed no effect in the assay, reconfirming the notion that it is probably not important. I received the original autoradiogram from the co-author who performed the Northern analysis in the 1988 study (Berleth et al., 1988), rescanned it once more to reveal the presence of both transcripts (Fig. 1A). *smbcd* is considerably less abundant but shows a much broader developmental expression profile than *lgbcd*, also being expressed during larval and pupal stages. Densitometric measurements indicate that in the 0-2 h lane, *lgbcd* is expressed roughly 200 times more strongly than *smbcd*.

*smbcd* arises through alternative splicing between exon 1 and 4, as illustrated in Fig. 1D. This splice event excludes exon 2 and 3, resulting in a small protein of 149 amino acids that lacks the homeodomain, as depicted in Fig. 1B. Intriguingly, this splice event appears to be evolutionary conserved. The reading frame to join exons 1 and 4 is conserved in many analyzed insect species, with confirmation obtained in *Drosophila melanogaster*, all Drosophilidae sequenced and *Lucilia sericata* to date. Notably, the splice event removes the carboxy-terminal part of a domain that possesses a regulatory function on the transcriptional activity of Lgbcd, termed the self-inhibitory domain (SID, Fig. 1B, C; (Zhao et al., 2003; Zhao et al., 2002). This domain was shown to have a tremendous effect on the transcriptional activity of Lgbcd. Mutations of 5 amino acids, either almost congruent to the last amino acids of exon 1 or the first amino acids of exon 2 (Fig. 1C), provoked dramatic changes in the transcriptional activity of Lgbcd, revealing a 25-fold increase when exon 1 amino acids were altered, or a nearly 70-fold decrease when amino acids from exon 2 were altered (Zhao et al., 2003). Since Smbcd results from of joining exon 1 and 4 (Fig. 1C), Smbcd has the potential to act as a transcriptional repressor for Lgbcd, mimicking a situation where amino acids of exon 2 were altered. Moreover, Lgbcd prefers to bind to other Bcd molecules when transcriptionally active, and the interaction was defined as being located NH2-terminally of the SID (Ma et al., 1996; Yuan et al., 1996). Hence, it is conceivable that Smbcd and Lgbcd can interact, most likely resulting in the downregulation of the transcriptional activity of Lgbcd. Unfortunately, *smbcd* was neglected and forgotten, and its role as a potential transcriptional regulator of Lgbcd was not discussed, nor was the congruence of the splice event and the altered transcriptional activity recognized (Zhao et al., 2003). Smbcd still harbors the PRD repeat (Frigerio et al., 1986) and the majority of an acidic domain (Fig. 1B).

Considerable efforts were made in the past to isolate *bcd* mutants, with a clear bias toward obtaining strong alleles (Frohnhöfer, 1987; Frohnhöfer and Nüsslein-Volhard, 1986; Seeger, 1989; Seeger and Kaufman, 1990), many of which turned out to be mutations around and within the homeodomain (Fig. 1D, red and orange asterisks). Notably, all classical *bcd* mutations are unlikely to affect *smbcd*. Another strong *bcd* mutant was generated by CRISPR-mediated insertion of a MiMIC cassette (Venken et al., 2011), inserting the cassette into the first intron and providing *trans*-splicing between exon 1 and the next exon on the MiMIC cassette (Huang et al., 2017). Again, this mutation is unlikely to affect *smbcd*. Three weak *bcd* mutations were recovered (Fig. 1D, green asterisks), with two showing lesions in the PEST domain, and another one showing a lesion in exon 4, common to both *lgbcd* and *smbcd* (Fig. 1D). The latter one, termed *bcd^2-13^*(Frohnhöfer and Nüsslein-Volhard, 1986; Struhl et al., 1989), now designated as *bcd^2^* by FlyBase (Attrill et al., 2016); Fig. 1D), when analyzed in *trans* to a *bcd* deficiency, revealed an almost complete wild-type cuticle pattern (Frohnhöfer, 1987; Frohnhöfer and Nüsslein-Volhard, 1987). Hence, the only allele also affecting *smbcd* was largely neglected as well.

Due to the apparent lack of a genetic situation where the complete *bcd* locus was deleted and to gain further insights into the function of *smbcd*, I opted to eliminate the complete function of *bcd* by deleting the entire locus using CRISPR. However, two major pitfalls exist for this approach. The first and less conspicuous one, is the existence of a *bcd* pseudogene, termed CR14578 (FlyBase; (Attrill et al., 2016; Frigerio et al., 1986), located in the heterochromatic region of 3L. This pseudogene shares almost 100% DNA sequence identity with over 2/3 of the *bcd* genomic region, including the homeodomain (Fig. 1D, green), but lacks a promoter, a transcription start site and upstream regulatory sequences. Furthermore, the entire exon 4 and the *bcd* 3’UTR are lacking, and no transcriptional activity was documented (Berleth, 1989). CR14578 was discovered in the homology screen that also allowed the isolation of the *bcd* gene (Frigerio et al., 1986). Hence, any attempt to derive gRNA sequences common to both genes would inevitably lead to low efficacy. The second pitfall is the erroneous opinion that the homeodomain located on exon 3 would be the prime target and sufficient for inactivation of the *bcd* gene, demonstrated in the past when strategies for a *bcd* knock-out were designed (Huang et al., 2017).

To address the problems described above, gRNAs were designed to target sequences outside of CR14578. This resulted in a deletion of approximately 3485bp within the *bcd* genomic region, spanning from the transcription start site to nearly the complete end of the 3’UTR (Fig. 1D, pink). Using homology-directed repair (HDR), the genomic region was then replaced by approximately 1.4 kb of sequences comprising an *eyeless* promoter controlling the expression of DsRed-Express, flanked by Lox-P sites (Rainbow Transgenic Flies Inc.), creating an allele termed *bcd^CRISPR^*. In another study (Fernandes et al., 2022), a similar deletion was introduced to create a second *bcd^CRISPR^* allele, termed *Δbcd^CRISPR^*. However, no in-depth analysis was conducted using this allele.

*bcd^CRISPR^* flies behaved identically as any of the classical *bcd* alleles, showing a strict maternal phenotype. Homozygous *bcd^CRISPR^* females exhibited a normal body size, but their abdomen was often wider than normal (data not shown). The main reason was that their ovarioles were somewhat shorter but wider, hence appearing round (Fig. 2A), as did a single egg (brown arrowhead, Fig. 2A). A mixture between *bcd^CRISPR^* embryos (Fig. 2B, brown arrowheads) and wild-type embryos (Fig. 2B, blue arrowheads) revealed the difference. This distinct geometry prompted me to measure the aspect ratio (length/width) of the two type of eggs as shown in Fig. 2C, measurements according to (Andersen and Horne-Badovinac, 2016). Whereas the value of the aspect ratio in control wild-type embryos centered around 2,38 (n=26; Fig. 2C, blue bars), that of *bcd^CRISPR^* embryos was considerably smaller, 1.83 (n=32), but showed more variation (Fig. 2C, brown bars). There is an important phenotypic distinction between round eggs, which have a decreased egg length and increased width, and short eggs, which only have a decrease in egg length. Both defects result in a reduced aspect ratio. In each of the independent 9 *bcd^CRISPR^* lines obtained from the mutagenesis experiment, egg length was significantly decreased, and egg width was increased (data not shown). These observations suggested that *bcd^CRISPR^* flies produce round eggs. *bcd^E1^/bcd^CRISPR^* transheterozygous or *bcd^E1^*/*bcd^E1^* homozygous eggs (the *bcd^E1^* allele was commonly used in most reports in the past, Fig. 1D), produced normally-sized embryos (Frohnhöfer, 1987; Frohnhöfer and Nüsslein-Volhard, 1986). Since *smbcd* is not affected in *bcd^E1^* mutants (Fig. 1D), there is a strong indication that the short and round egg phenotype is primarily due to *smbcd* and not due to *lgbcd*. Consistent with this observation is that a *bcd^2-13^* embryo, mentioned above, and as shown in (Nusslein-Volhard et al., 1987) is substantially smaller than the other embryos of the *bcd* phenotypic series. Finally, the notion that the small egg phenotype is primarily due to *smbcd* is supported by analyses of *smbcd* mutant eggs which also reveal small and round eggs (S. B., in preparation).

**Figure 2.**
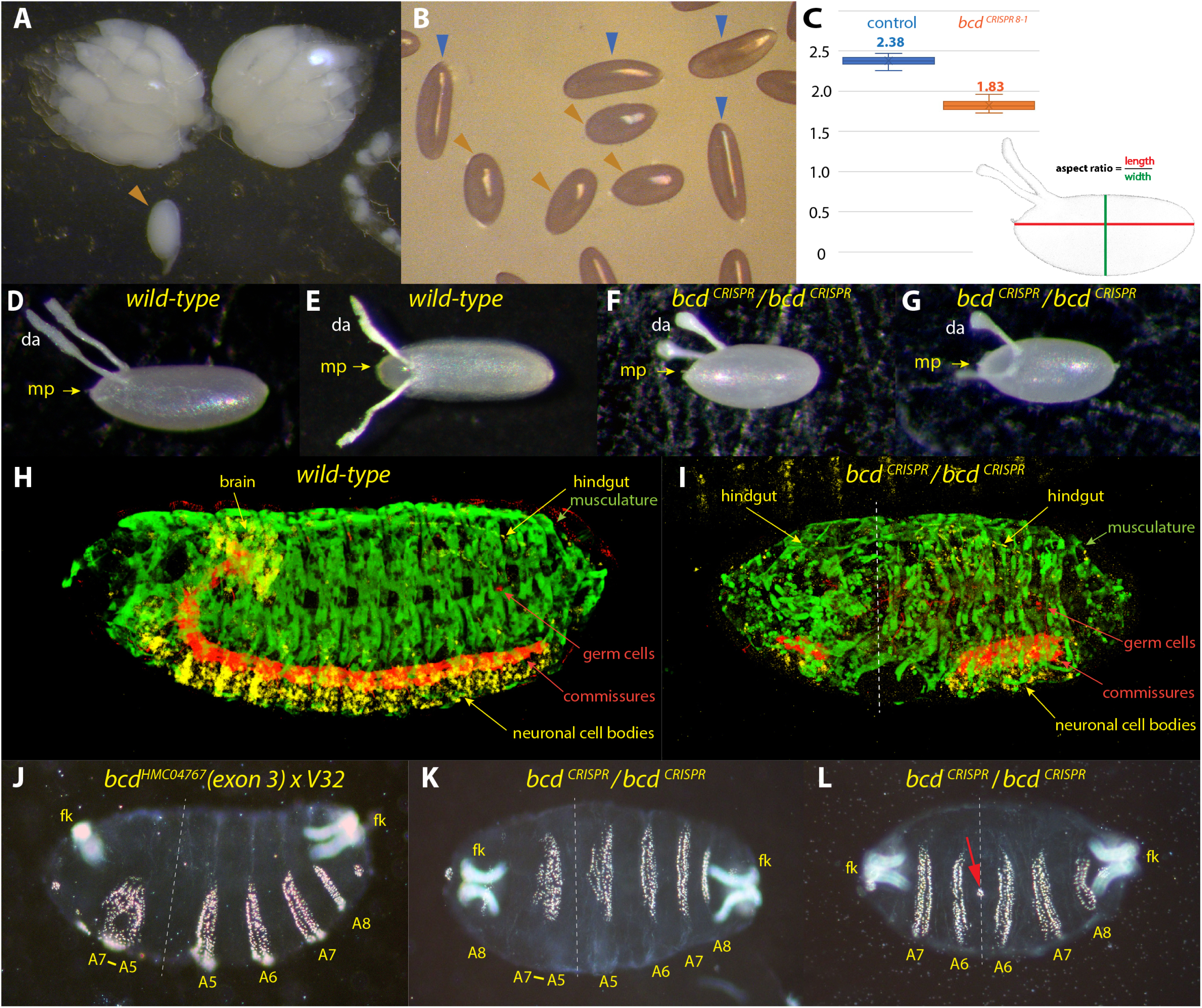
*bcd* is required for the shape of the egg. **(A)** dissected ovarioles of *bcd^CRISPR^* mothers, showcasing an overall round appearance, along with a single late-staged oocyte (brown arrowhead). (B) egg collection displaying a mixture of wild-type eggs (blue arrowheads) and *bcd^CRISPR^* eggs (brown arrowheads). (C) aspect ratio calculated as length/width, as outlined in the figure of dechorionated control embryos (blue bars, n=26, mean 2.38) and dechorionated *bcd^CRISPR^* embryos (brown bars, n=32, mean 1.83), with measurements following (Andersen and Horne-Badovinac, 2016). (D, E) wild-type eggs, presented in lateral (D) and dorsal (E) views to illustrate the morphology of dorsal appendages. (F, G) *bcd^CRISPR^* eggs, shown in lateral (F) and dorsal (G) views, revealing shorter but thicker dorsal appendages. (H-I) embryonic stage 15 3D-embryos from a confocal stack using different protein stainings and colors to stain various tissues: β3-Tubulin (green) stains muscles; Runt (yellow) stains neuronal nuclei of the CNS and brain and the hindgut (yellow arrow), a combination of mab BP102 and Vasa (red) stains the ventral chord and germ cell (GC), respectively, of a wild-type embryo in (H) and a *bcd^CRISPR^* embryo in (I). Note the irregular duplicated musculature at the anterior and the interrupted CNS in (I). Also observe the asymmetry of the embryo regarding the duplicated posterior end (dashed line). (J) cuticle phenotype from a cross of a *bcd* RNAi line, *bcd^HMC04767^* x *V32* (maternal driver) creating a stronger phenotype than any of the “classical” *bcd* alleles (Frohnhöfer and Nüsslein-Volhard, 1986, 1987). (K) *bcd^CRISPR^* cuticle exhibiting a rounder shape and a phenotype even stronger than that in (J). (L) extreme, but less frequent *bcd^CRISPR^* phenotype. Note the short and round phenotype as a result of the superimposed egg geometry and embryonic patterning phenotype of complete loss of *bcd*. The red arrow points towards a small circle of denticles infrequently seen in some embryos, see also supplemental Fig. 1G. Dashed lines in (I-L) denote planes of symmetry for duplication of posterior tissues to the anterior side, best seen as duplications of the filzkörper (fk) as a marker for posterior identity. All embryos are oriented anterior to the left and dorsal side up, unless otherwise noted.

Intimately linked to the egg chamber are the eggshell structures that house the egg chamber and later also the embryo. A prominent structure on the dorsal side are the dorsal appendages (da) that arise from specialized dorsal follicle cells (Pyrowolakis et al., 2017); Fig. 2D, E). These structures appear as long fine threads. In *bcd^CRISPR^* mutants, the appendages were much shorter but thicker (Fig. 2A, F, G), akin to changes of egg geometry in the mutants. However, the position and shape of the micropyle, another structure of the eggshell (Horne-Badovinac, 2020) remained unchanged in *bcd^CRISPR^* mutants (Fig. 2D, F, G; mp, yellow arrows).

Prompted by the changed egg geometry in *bcd^CRISPR^* mutants, I reasoned that the layout of the different germ layers in the *bcd^CRISPR^* mutant embryo would be changed. To investigate this, I stained wild-type and mutant eggs with a panel of antibodies to determine the fate of the different tissues. 3-D reconstructions of embryos, stained with important tissue markers and recorded by widespread confocal stacks, were produced. This allowed monitoring the spatial changes of the different tissues in the mutant. The mutant embryo (Fig. 2I) exhibited several defects. The first one concerned a clear asymmetry regarding the size of the duplicated posterior tissues (dashed line). There was an apparent disorganization of the musculature (green), to a lesser extent in the authentic posterior tissue, but pronounced in the duplicated tissues, suggesting that the tissue patterning was not determined properly. Neuronal tissues were induced (yellow, red), but were interrupted at the mirror plane, evident by the ruptured commissures. Endodermal tissues, such as the hindgut (yellow), appeared duplicated in the correct anterior-posterior direction. Lastly, the germ cells (in red) were absent in the duplicated posterior part. Regarding the scaling of internal organ to match the size of artificially size-reduced embryos (Tiwari et al., 2021), comparable organs, such as the hindgut or neuronal tissue, exhibited scaling in the duplicated anterior tissues.

While tissue analysis of faulty tissue organization allowed for a crude allocation of the defects in the mutant embryo, the analysis of the cuticle pattern served as a precise tool for determining how a gene influences the patterning of the anterior-posterior axis of the embryo. As a reference for the *bcd* mutant, a *bcd* RNA^i^ line (*bcd^HMC04767^*) with a dsRNA target in exon 3 specific for *lgbcd*, as well as the *bcd* pseudogene CR14578, was used. When *bcd^HMC04767^* was driven by a strong maternal driver *V32*, it exhibited a strong *bcd* phenotype (Fig. 2J), with abdominal segments A8-A5 present and appearing normal. Anteriorly, fused abdominal segments A7-A5 and the filzkörper (fk) were duplicated and arranged mirror-invertedly, but the latter was not fully developed. Notably, A8 was missing and therefore, not duplicated, creating an asymmetry with respect to the axis of reflection. These results were consistent with previous analyses on *bcd* cuticles using various *bcd* RNA^i^ lines and varying the amount of GAL4 activity (Staller et al., 2015).

In *bcd^CRISPR^* mutants, a wide range of cuticle phenotypes was observed (Fig. S1; (Frohnhöfer, 1987; Frohnhöfer and Nüsslein-Volhard, 1986; Huang et al., 2020; Staller et al., 2015), with the phenotype in Fig. 2K representing the most frequent pattern defect. In rare cases, a weak anterior patterning defect was observed (Fig. S1D), along with an increasing frequency of stronger anterior patterning defects (Fig. S1E-G). Embryos of the most frequent pattern defect were round and exhibited A8-A5 on the posterior side (Fig. 2K), but the content of the duplicated posterior end was not identical to that in the RNA^i^ line (Fig. 2J); instead, it revealed A8 as an independent unit, with only A7-A5 fused. In rarer cases of *bcd^CRISPR^* cuticles, very round embryos were visible with only A8-A6 apparent and appearing duplicated anteriorly (Fig. 2L). Interestingly, in more severe *bcd^CRISPR^* cuticles, the formation of the duplicated anterior filzkörper was complete, in contrast to that of the *bcd^HMC04767^* allele (Fig. 2J) or the weaker phenotypes (Fig. S1D-F). The completeness appeared to depend on the geometry of the embryo, as previously noted (Frohnhöfer, 1987; Huang et al., 2020). It is worth noting that anterior cuticle defects were occasionally discovered in larvae from heterozygous *bcd^CRISPR^* females, where T1 and T2 patterning was poorly manifested or largely absent (Fig. S1B, C). In conclusion, the complete removal of both *smbcd* and *lgbcd* had a more significant impact on the shape of the embryo and resulted in more severe pattern defects than the removal of *lgbcd* alone. However, both genetic backgrounds still exhibited a wide variation of phenotypes.

Preliminary analysis of *smbcd* expression revealed that there is robust expression in the germinal vesicle (GV) (S. B., in preparation). Historically, only a few proteins have been described to be expressed in the GV, with the prevailing belief that the GV is barely transcriptionally active, except during oocyte stage 10a (Mahowald and Tiefert, 1970). To explore this further, I conducted a screening for the expression of patterning genes in the GV, focusing on candidate genes for interaction with *bcd* based on interaction data from the embryo (Attrill et al., 2016). The first significant hit was *hunchback* (*hb*) (Fig. 3A), revealing expression not only in the nurse cell nuclei but also in the GV. Notably, this expression has not been reported previously. Again, Hb expression in the oocyte was often overlooked in the past, due to its established importance for segmentation in the embryo. Next, *lesswright* (*lwr*), a gene implicated in Lgbcd import into the nuclei during early embryogenesis (Epps and Tanda, 1998) showed a similar GV localization (Fig. 3B), except for being excluded what appeared to be the Cajal body (Fig. 3B, insert). The Cajal body is a subnuclear organelle involved in various cellular processes, and its exclusion in this context may suggest specific regulatory mechanisms. Finally, *maelstrom* (*mael*), a gene of the *spindle* (*spn*) class of segmentation genes, which produces eggshells with variable anteroposterior and dorsoventral axis defects (Schupbach and Wieschaus, 1991) showed a prominent GV expression (Fig. 3C).

**Figure 3.**
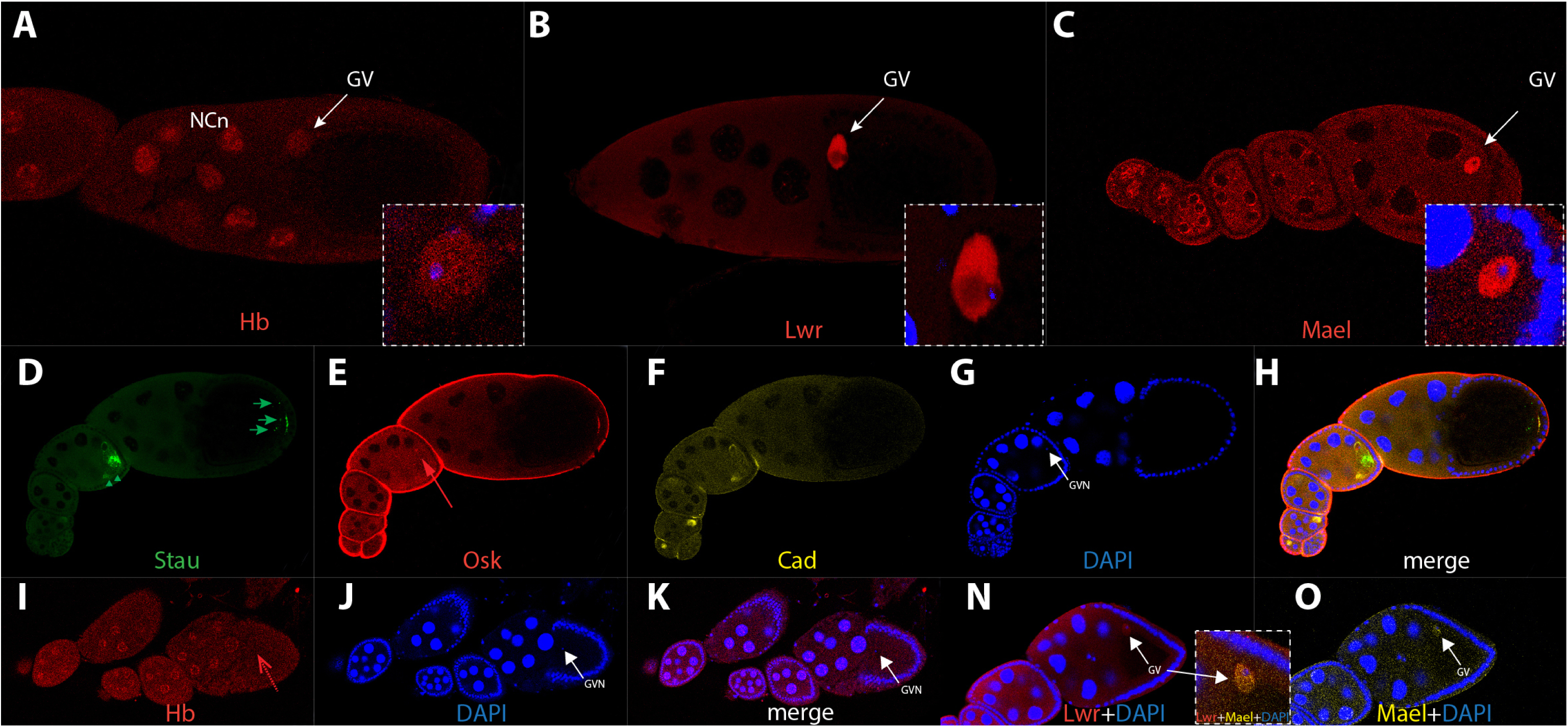
Expression of axis-patterning genes in wild-type and *bcd^CRISPR^* oocytes. (A) wild-type stage 10 oocyte stained for Hunchback (Hb); both the nurse cell nuclei (NCn) and the germinal vesicle (GV, arrow) exhibit Hb staining. Insert displays a magnification of the GV staining (red) along with DAPI (blue). (B) wild-type stage 10 oocyte stained for Lesswright (Lwr); the position of the germinal vesicle (GV) is indicated with an arrow. Insert features a magnification of the GV staining (red) along with DAPI (blue). (C) wild-type stage 10 oocyte stained for Maelstrom (Mael); the position of the germinal vesicle (GV) is indicated with an arrow. Insert shows a magnification of the GV staining (red) along with DAPI (blue). (D-H) *bcd^CRISPR^* ovariole stained for Staufen (D, green), Oskar (E, red), Caudal (F, yellow), DAPI (G, blue) and merge in (H). Posteriorly, Staufen shows partial mislocalization (green arrowheads and arrows in D). Oskar staining at the apices of follicle cells is unspecific staining of the mouse anti Osk antibody. (I-K) *bcd^CRISPR^* oocyte string stained for Hb (I, red), DAPI (J, blue) and merge in (K). Hb staining is no longer detectable in the GV (stippled red arrow in I, arrows in GV nucleus (GVN) in J and K), while the nurse cells nuclei still express Hb. (N, O) *bcd^CRISPR^* stage 8 oocyte, Lwr (N) and Mael (O) expressions are unaffected in the GV (arrows). The insert between (N) and (O) shows a merge of Lwr, Mael and DAPI staining. All oocytes are oriented anterior to the left. Stages of oogenesis follow the classification of (King, 1970).

Several decades ago, translation of *lgbcd* was claimed to not occur during oogenesis (Driever and Nüsslein-Volhard, 1988), recently refuted by (Ali-Murthy and Kornberg, 2016), who showed Lgbcd expression during late stages of oogenesis at the anterior rim of the oocyte, in a pattern reminiscent of that of the mRNA. Hence, since in the past control of other maternal genes by *bcd* was considered unlikely, thorough analyses on the role of *bcd* on other maternal genes were not conducted. To correct this picture and to define the function of *bcd* during oogenesis, I looked at the expression pattern of some anterior-posterior patterning genes in *bcd^CRISPR^* mutants. The first one was *staufen* (*stau*) which revealed a slightly disorganized protein localization at early stages (Fig. 3D, arrowheads), while during later stages, posterior localization was incomplete with particles remaining in the bulk of the oocyte (Fig. 3D, arrows). *oskar* (*osk*) during early stages showed a distinct protein localization adjacent to the GV (Fig. 3E, arrow); however, its later posterior localization at the polar plasm was not affected, suggesting that complete removal of the *bcd* locus did not affect the MT-based localization process. *caudal* (*cad*), another segmentation gene showed strong expression in wild-type oocytes (data not shown), although its expression profile was sparsely documented during this stage (Mlodzik and Gehring, 1987a) and no maternal Cad protein localization was reported. In *bcd^CRISPR^* mutants, Cad localization appeared unchanged during all stages, however, its expression levels were dramatically increased during early stages (Fig. 3F), consistent and in analogy to the documented translational repression of *cad* by Bcd in the embryo (Niessing et al., 2002); Fig. 4).

**Figure 4.**
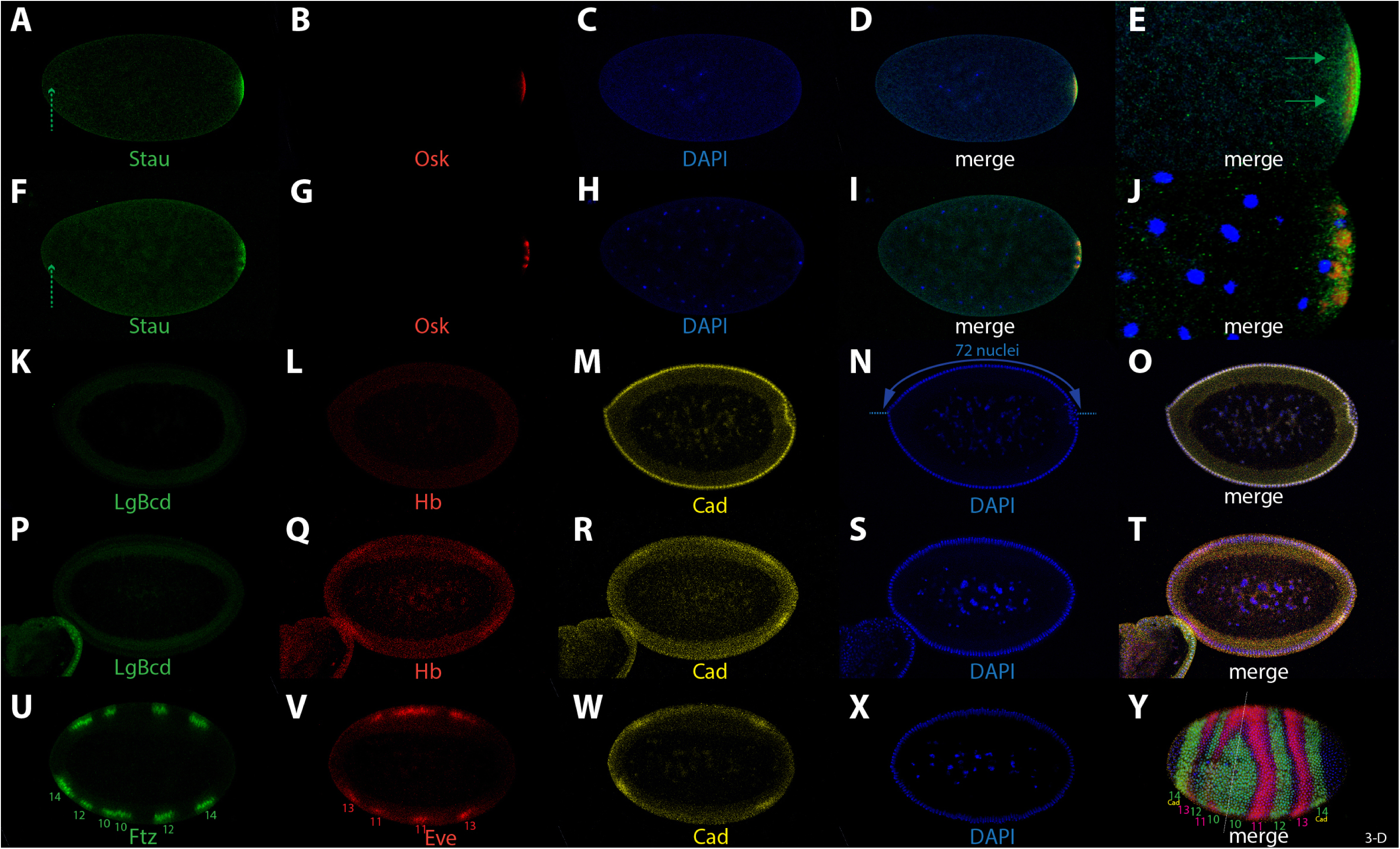
Influence of complete absence of *bcd* on segmentation genes in the embryo. (A-E) *bcd^CRISPR^* nc 4 embryo stained for Staufen (A, green), Oskar (B, red), DAPI (C, blue) and merge in (D). (E) high magnification of the posterior end after 3-D reconstitution of the whole stack. Staufen is not localized at the anterior tip due to absence of *bcd* mRNA (A, stippled green arrow) and appears loosely associated at the posterior end (E, green arrows). (F-J) *bcd^CRISPR^* nc 10 embryo stained for Staufen (F, green), Oskar (G, red), DAPI (H, blue) and merge in (I). (J) high magnification of the posterior end after 3-D reconstitution of the whole stack. Staufen is not localized at the anterior tip due to absence of *bcd* mRNA (F, stippled green arrow). Osk is no longer uniformly localized in the posterior polar plasm (PP) but appears clustered in distinct domains within the PP. (K-O) *bcd^CRISPR^* early nc 14 embryo stained for Lgbcd (K, green), Hb (L, red), Cad (M, yellow), DAPI (N, blue) and merge in (O). Hb is not activated due to absence of *bcd*, and Cad is expressed uniformly along the A-P axis due to lack of repression by *bcd*. Note the reduced number of blastoderm nuclei, annotated in (N). (P-T) *bcd^CRISPR^* late nc 14 embryo stained for Lgbcd (P, green), Hb (Q, red), Cad (R, yellow), DAPI (S, blue) and merge in (T). The late posterior Hb (Q) and Cad (R) bands appear activated at the anterior due to adoption of posterior identities at the anterior. (U-Y) *bcd^CRISPR^* late nc 14 embryo stained for Ftz (U, green), Eve (V, red), Cad (W, yellow), DAPI (X, blue) and merge in (Y) as a 3-D reconstruction of the whole stack. Numbers in (U, V and Y) indicate numbering of parasegments (PS) in their respective color of staining. The dashed line in (Y) denotes the mirror plane for the duplication of posterior tissues to the anterior. All embryos are anterior to the left and dorsal side up, unless otherwise noted. Note: due to the mounting procedure, embryos are flattened and may appear somewhat rounder than they actually are.

I then asked if the peculiar GV stainings of patterning genes would be dependent on *bcd*. To this end, I analyzed their patterns in *bcd^CRISPR^* oocytes. Hb, a direct target of *bcd* in the embryo (Driever and Nusslein-Volhard, 1989; Tautz, 1988), revealed absence in GV staining (Fig. 3I, stippled arrow), while expression in the nurse cell nuclei was unaffected, suggesting a distinct regulation of processes in the GV compared to those in the NCs. Lwr and Mael expression in the GV, on the other hand, were unaffected in *bcd^CRISPR^* mutant oocytes (Fig. 3N, O), also with respect to subnuclear localization within the GV.

As for the dependence of prominent segmentation genes on *bcd* in the embryo, most of them relied on *lgbcd* activity (Fig. 4), as could be expected based on previous analyses using the classical *bcd* alleles (Liu et al., 2013; Staller et al., 2015). However, the geometry of *bcd^CRISPR^* eggs changed dramatically compared to that of the relatively normal “classical” *bcd* alleles (Frohnhöfer, 1987; Frohnhöfer and Nüsslein-Volhard, 1986), introducing a new constraint: due to the reduced volume of the egg, not all of the approximately 6000 nuclei that normally populate the wild-type blastoderm embryo (Zalokar and Erk, 1976) found space at the cortex. Instead, a considerably lower number of nuclei assembled at the cortex. As evident in Fig. 4N, the *bcd^CRISPR^* blastoderm embryo could host only around 70-75 nuclei along the dorsal side, compared to 106 ± 3 nuclei in wild-type embryos (Suksuwan et al., 2017). My method of measuring the number of nuclei differed from that described in (Huang et al., 2020). I measured from the middle of the anterior tip to the middle of the posterior tip in single mid-sagittal confocal sections (Fig. 4N; (Cai et al., 2017; Spirov et al., 2009), hence the number in this study is larger than that of (Huang et al., 2020). Of note, the interior yolk did not contain any excess nuclei that could not move to the periphery during cortical migration (Baker et al., 1993), nor was there a double layer of nuclei present at the cortex. I can conclude that in the *bcd^CRISPR^* mutant background, there is a system in play that adapts the final number of nuclei to the size of the embryo. Whether this step involves modulating the activity of nuclear divisions or degrading existing nuclei is currently unclear.

Since Stau localization in the oocyte was compromised in *bcd^CRISPR^* mutants, a similar behavior in the early embryo was anticipated. Indeed, posterior accumulation was compromised as well (Fig. 4A, E), indicating that not all Stau protein accumulated at the posterior rim, which later becomes the polar plasm. Moreover, since the *bcd* mRNA was crucial for Stau localization to the anterior pole (Ferrandon et al., 1994), Stau did not localize anteriorly in a *bcd^CRISPR^* embryo at all (Fig. 4A, F, stippled arrow). On the other hand, posterior Osk localization in the polar plasm was unaffected (Fig. 4B), as observed in the oocyte (Fig. 3E). In later staged *bcd^CRISPR^* embryos, e.g., at syncytial blastoderm during nc 10, aberrant posterior Stau localization became more prominent and showed a loosely dispersed appearance at the posterior pole (Fig. 4F, J). Again, Osk showed a normal localization and even became normally internalized into the pole cells (Fig. 4G, J).

Next, I tested the dependence of known segmentation genes in the embryo on the complete absence of *bcd*. A prime target was *hb* whose expression was no longer visible, and the broad anterior band disappeared (Fig. 4L), as did Lgbcd (Fig. 4K). Hence, *hb* behaved quite differently in *bcd^CRISPR^* embryos compared to the pattern in the classical *bcd* alleles where low-level Hb staining was observed (Tautz, 1988). The interpretation at that time was that *lgbcd* would be critically required for the onset of *hb*, while the weak observed *hb* expression was a consequence of residual maternal *hb* (Tautz, 1988). Cad, on the other hand, behaved as expected from the strong *bcd* classical alleles (Mlodzik and Gehring, 1987a; Mlodzik and Gehring, 1987b; Niessing et al., 1999). Instead of the shallow posterior-to-anterior gradient derived from the maternal *cad* transcript (Macdonald and Struhl, 1986; Mlodzik and Gehring, 1987a), strong ubiquitous expression along the entire anterior-posterior axis was observed (Fig. 4M), suggesting complete derepression of translational control of Cad by *bcd* (Niessing et al., 2002; Niessing et al., 1999).

At late blastoderm stage (nc14), in wild-type embryos, apart from the broad anterior band, a smaller second posterior band of Hb became visible (Tautz, 1988). In *bcd^CRISPR^* mutants, this band was present and appeared duplicated at the anterior (Fig. 4Q), as it did in the strong classical *bcd* alleles, demonstrating that the anterior tissues had adopted posterior identity. For Cad, in late nc 14 wild-type embryos, the shallow posterior to anterior gradient disappeared, and, driven by the zygotic *cad*-promoter, a new posterior Cad band showed up (Macdonald and Struhl, 1986; Mlodzik and Gehring, 1987b). In *bcd^CRISPR^* mutants, the late posterior Cad domain appeared duplicated at the anterior (Fig. 4R, W), as could be expected from data of the strong classical *bcd* alleles (Mlodzik and Gehring, 1987b). Kr expression in *bcd^CRISPR^* mutants was not detectable (data not shown), suggesting that the segmental anlagen where Kr is expressed in wild-type embryos was absent or not defined in the mutants.

I then investigated the impact of *bcd* on the next class of the hierarchy of segmentation genes, the pair-rule genes, focusing on *fushi-tarazu* (*ftz*) and *even-skipped* (*eve)*, which define even-numbered and odd-numbered parasegments, respectively (PS; (Lawrence et al., 1987). The key question was how the segmental axis would behave in an egg with dramatically altered geometry and a reduced number of nuclei. In wild-type embryos, both pair-rule genes displayed seven regularly-spaced stripes, each 3-4 cells wide and showing reciprocal patterns of expression (Frasch and Levine, 1987). In *bcd^CRISPR^* mutants, only the posterior tissues became specified and appeared duplicated at the anterior (Fig. 2K, L). Consequently, the identity of the pair-rule stripes in a *bcd^CRISPR^* mutant reflected this behavior. The Ftz pattern exhibited a reduced set of stripes, with posterior stripes of PS 14, 12 and 10 present, along with the same set added inverted at the anterior side (Fig. 4U). Similarly, Eve showed the posterior PS 13 and 11 stripes, along with the same set inverted at the anterior pole (Fig. 4V). The posterior Cad domain aligned with that of the posterior-most Ftz stripe, i. e., PS 14 (Fig. 4W, Y) and in the duplicated anterior tissue, congruence of both expression domains was maintained.

In a 3-D analysis (Fig. 4Y), it became evident that the process of duplication of posterior tissue was not symmetrical, not equivalent in size, and not constant along the dorsal-ventral (D-V) axis. The mirror plane (Fig. 4Y, stippled line) was shifted anteriorly dividing the embryo into two halves with a width ratio of 37% vs. 63%. The developmental program then converted this asymmetry into a larva where the ratio was less pronounced but still recognizable (Fig. 2E, F). This plasticity likely occurred through embryonic pattern repair mechanisms involving apoptosis and changes in mitotic domains along the anterior-posterior axis, as shown in embryos with a triple dose of *bcd* (Namba et al., 1997). Another observation was that the numbers of nuclei of the Ftz and Eve bands on the authentic posterior side were abnormally high, exceeding those in wild-type embryos. The layout and the molecular determinants for the parasegments were altered. In the posterior part, PS 14 and 12 Ftz stripes as well as PS 13 and 11 Eve stripes measured 5-6 cells instead of 3-4 cells. Few of the apparent molecular changes at blastoderm were implemented in the remnant abdominal larval cuticle (Fig. 2E, F), again likely due to tissue repair (Namba et al., 1997). On the duplicated anterior part, the opposite was observed: too few nuclei, in the range of 2-3, expressed Ftz and Eve, with the exception of the anterior-most band of Ftz (PS 14) which appeared normal compared to wild-type (Carroll and Scott, 1985; Krause et al., 1988). A third noteworthy observation was the discrepancy of expression along the D-V axis of some bands. This was particularly evident near the axis of reflection (stippled line in Fig. 4Y). On the dorsal side, the PS 10 Ftz band was suppressed, while appears expanded on the ventral side. Likewise, PS 11 Eve band appeared fused on the dorsal side, but its anterior duplicate was not activated on the ventral side. Hence, this data suggested that *bcd* might even have some implications on the determination of the D-V axis.

All *bcd^CRISPR^* mutant embryos corresponded to range 1 and 2 embryos in the analysis by (Huang et al., 2020), where embryonic geometry was artificially altered through downregulation of the *fat2* gene during oogenesis, resulting in round embryos (Horne-Badovinac et al., 2012). The analysis of (Huang et al., 2020) was performed in a *bcd^MiMIC^* mutant background (Fig. 1D), which unfortunately still retains *smbcd^+^* activity. All segmentation gene patterns analyzed in this report (Kr, Hb, Gt (not shown), Eve, Ftz) in *bcd^CRISPR^* mutant embryos conformed to those patterns, suggesting that the genetic condition of *bcd^CRISPR^* alone was sufficient to provide conditions for the analysis under decanalized conditions.

## Discussion

For over three decades, the scientific community has relied on a phenotype of an important *Drosophila* patterning gene, *bcd*, encompassing a plethora of alleles, with the strong ones widely believed to be *null* alleles. This gene has served as a paradigm for morphogenetic gradients in biology textbooks. However, the incomplete description of the phenotype can largely be attributed to the inadequate reproduction of the Northern analysis in (Berleth et al., 1988) and the striking appearance of the Lgbcd gradient (Driever and Nüsslein-Volhard, 1988). The gradient confirmed earlier predictions by (Frohnhöfer and Nüsslein-Volhard, 1986) regarding a morphogenetic substance, as established through genetic studies and cytoplasmatic transplantation experiments. Consequently, everyone was satisfied, and the quest for the function of the other *bcd* isoforms became superfluous, leading to complete oblivion of *smbcd*.

In this paper, I have demonstrated a new and fundamental feature and function for *bcd*: its influence on egg geometry. The change in the geometry of *bcd^CRISPR^* eggs prompts the question: What causes this change? Significant progress has been made in understanding how the egg is shaped during oogenesis and how the egg chamber structure and morphogenesis occur (Bilder and Haigo, 2012; Cetera and Horne-Badovinac, 2015; Osterfield et al., 2017). A crucial aspect of these processes is the interplay during the growth of the initially round follicle (egg chamber) with the surrounding muscle sheath and the basement membrane (BM), which help shape the geometry of the egg chamber and to elongate it during growth, collectively referred to as the molecular corset (Andersen and Horne-Badovinac, 2016). A recent report showed that tissue elongation is controlled by a single pair of cells at the anterior pole, the pole cells (PCs), which secrete matrix metalloproteinase (MMP) that senses the degree of tissue elongation (Ku et al., 2023). It was concluded that when the anterior PCs increase the levels of MMP1, the thickness of the BM fibers increases, and the corset becomes more resistant to pressure exerted from the growing follicle, leading to expansion along the anterior-posterior axis. Since control is exerted from the anterior pole, only *lgbcd* would be in place at the anterior rim of the oocyte (Ali-Murthy and Kornberg, 2016) to regulate the expression of MMP1 in the PCs. However, classical *lgbcd* mutant embryos exhibit a rather normal geometry (Frohnhöfer, 1987). Nevertheless, since there is another pair of PCs at the posterior pole, these PCs may represent another control center for tissue elongation.

It is apparent that the shape of the embryos depends on *smbcd* inputs during oocyte growth. Absence of *smbcd* only results in short and round embryos (S. B., in preparation). Conversely, in the other extreme case where excess *smbcd* is provided, as seen in *bcd^+5+8^* embryos (Ali-Murthy and Kornberg, 2016; Cai et al., 2017; Frohnhöfer and Nüsslein-Volhard, 1986; Namba et al., 1997), the embryo shape changes, showing a slight increase in both length and width, typically with a pointed anterior end. The aspect ratio often exceeded that of wild-type embryos (Cai et al., 2017). It can thus be speculated that the more copies of *smbcd* during oogenesis are present, the longer and wider the embryo might become. Whether this step requires only *smbcd* or *smbcd* in conjunction with *lgbcd* is currently unclear. Both isoforms are translated during oogenesis, suggesting their likely action at the protein level (Ali-Murthy and Kornberg, 2016); S. B., in preparation).

Regarding *bcd* being a master regulator of patterning processes, a recent report demonstrated that *hb* activates *bcd* in Pair1 neurons, rather than *vice-versa* (Feng, 2022; Lee et al., 2022). This finding challenges the traditional view that key players in embryonic patterning processes maintain fixed roles. Moreover, it indicates a novel role for *bcd* beyond embryonic patterning, challenging our belief that *bcd* is only responsible for the patterning processes during the first 14 nuclear cycles in the embryo. Northern analysis confirms a more widespread expression (Fig. 1A). Sparse and punctated expression of *bcd* in Pair1 neurons based on liquid-liquid phase separation (LLPS, (Liu et al., 2020)) may often escape our detection beyond the first 3 hours of development. In the case of oocyte patterning, *hb* again is clearly regulated by *bcd* in its activity in the GV but does not appear to be dependent on *bcd* in its expression in the NCn (Fig. 3I).

The splice event leading to *smbcd*, involving the joining of exon 1 to exon 4 and following a phase zero behavior, appears to be a highly conserved step during Cyclorrhapha evolution, the sub-order where *bcd* is present in the genomes. This phase zero splice event correlates with the structure of ancient proteins (de Souza et al., 1998). Within this scenario, a conserved alternative splicing event exhibits increased selection pressure to preserve the frame (Resch et al., 2004). Consistently, sequences in intron 1 and 3 (flanking exon 1 and 4, 559 bp and 513 bp in size, respectively) show several stretches of elevated sequence identities up to 97% among related Drosophilidae. Therefore, a likely conserved machinery ensures correct splicing and adjustment of the correct ratio between *lgbcd* and *smbcd*.

In the past, *eGFP-bcd* genomic constructs were used to demonstrate the full functionality of the construct. If successful, it was often stated that …”the construct fully rescued a *bcd* phenotype”. Despite the compelling notion of a “full rescue”, relying solely on such a statement is deemed unfair. This is because it does not delineate the defects that may arise when *bcd* is active. Rather, it is the cumulative impact of defects at the blastoderm stage and the extensive repair mechanisms during subsequent developmental stages, as elucidated by (Namba et al., 1997) that eventually restore the defects back to wild-type state. Therefore, in the interest of fairness, researchers are encouraged to meticulously document any defects that manifest already at the blastoderm stage.

Notably, classical *bcd* mutants, previously thought to eliminate *bcd* function, are far from being *bcd^null^* mutants. This realization immediately poses two major problems: 1) it challenges the interpretation of past conclusions drawn from *bcd* rescue constructs in a *bcd* mutant background. These constructs often lead to an imbalance in *bcd* function, rather than to a return to a wild-type situation. For instance, flies with a rescue construct now have four copies of *smbcd* under such conditions, likely tipping the balance away from wild-type. 2) Numerous measurements of the diffusion constant done with eGFP-Bcd in the past (Abu-Arish et al., 2010; Athilingam et al., 2023; Drocco et al., 2011; Drocco et al., 2012; Durrieu et al., 2018; Gregor et al., 2007a; Gregor et al., 2007b; Grimm and Wieschaus, 2010; Huang et al., 2017; Huang et al., 2020; Little et al., 2011; Mir et al., 2018; Singh et al., 2022) can no longer be interpreted as resulting solely from Lgbcd protein. The signal likely comprises a mixture of the output of two copies of *lgbcd* and four copies of *smbcd*. Since the size of the protein determines its diffusion, the contribution of Smbcd has the strong potential to perturb any eGFP-Bcd read-out. Thus, conclusions drawn regarding the diffusion constant of Lgbcd are likely prone to become meaningless.

Much work lies ahead to obtain validated values of the true diffusion constants of Bcd isoforms. Most importantly, those of the Lgbcd and Smbcd alone, and not that of the mixture.

## Conclusions

This report unveils a new function of *bcd* in shaping egg geometry. The discovery of this novel *bcd* function requires a comprehensive revision of our understanding of this important patterning gene. Furthermore, past experiments conducted under the assumption that *bcd* mutants were *bcd^null^* mutants, which they were not, must be reevaluated.

## List of abbreviations

bcd: bicoid
lgbcd: large bicoid
smbcd: small bicoid
nos: nanos
stau: staufen
osk: oskar
lwr: less-wright
mael: maelstrom
lamC: lamin C
cad: caudal
hb: hunchback
eve: even-skipped
ftz: fushi tarazu
MMP: matrix metalloproteinase
gRNA: guide RNA
FISH: fluorescent in situ hybridization
ARTS: active mRNA transport, synthesis
SDD: synthesis, diffusion, degradation
MT: microtubule
A-P: anterior-posterior
RNP: ribonucleoprotein
aa: amino acid

## Declarations

### Ethics approval and consent to participate

not applicable

### Consent for publication

not applicable

### Availability of data and material

The datasets and material generated during and/or analyzed during the current study are available from the author on reasonable request.

### Author’s contribution

S. B. was responsible for conception, production and analysis of the data of Figs. 1-4 and S1, and for writing of the manuscript.

## Acknowledgements

I thank the Swedish Research Council, the Ekhaga -, Pia Ståhl -, Nilsson Ehle - and the Erik Philip-Sörensen-Foundation for support. I wish to thank Thomas Mayer for hosting me during a sabbatical stay in 2018 and subsequent years, Patrick Müller for continued support in his lab from 2022 on, and the Konstanz Bioimaging Facility for excellent maintenance of their LSM 700 confocal microscope. I thank Daniel Bopp for providing the original *bcd* Northern filter from 1986 and Markus Noll for permission to publish it once more. I also would like to thank Sol DaRocha for excellent and reliable technical assistance.

## Materials and methods

### *Drosophila* stocks and genetics

Complete *bcd* gene removal and subsequent replacement of the gap by a DsRed-Express reporter cassette under *eyeless* control was designed and fabricated by Rainbow Transgenic Flies Inc. using proprietary standard technology. Special attention was given to designing the guide RNAs to ensure that they do not overlap with sequences of pseudogene CR14578, which shares 100% identity with *bcd* over a larger stretch. Such an overlap could potentially affect the efficacy of the strategy. gRNA1 target: GATCGCAAAAACGCAAAATG-TGG, at the transcription start site; gRNA2 target (lower strand), GCTTAAAGAGACAACATCAA-AGG, about 40 nucleotides upstream of the poly A site. This approach results in the deletion of approximately 3485 base pairs from the *bcd* locus leading to absence of both *lgbcd* and *smbcd*, as confirmed by PCR and antibody staining for Lgbcd (Fig. 4P) and Smbcd (data not shown). The integration of the reporter gene within the *bcd* locus was verified through PCR and subsequently confirmed by sequencing. Individual fly lines were balanced using the balancer chromosome *TM3 Sb*, termed *bcd^CRISPR^* and exhibited the expected behavior, showing a strict maternal phenotype as do the strong classical *bcd* alleles.

Egg morphology (aspect ratio length/width) analyses were done according to (Andersen and Horne-Badovinac, 2016), using dechorionated embryos through a standard bleaching protocol.

The *bcd* RNA^i^ line in Fig. 2 was Bloomington stock number 57458 with the genotype *y1 sc* v1 sev21; P[TRiP.HMC0476] attP40* targeting a sequence in exon 3, hence is specific for *lgbcd*. However, this strain also targets the *bcd* pseudogene CR14578. The strong maternal V32 driver was used, and crosses were exposed to 29° for maximal expression of the transgene. Flies were fed with standard fly food (Bloomington recipe).

### Cuticle preparations

Embryos were collected in 24 hrs. interval, incubated > 36 hrs., dechlorinated in 50% bleaching solution, fixed in 25% formaldehyde for > 5 hrs., devitellinized, mounted in Hoyer’s medium and incubated at 65^°^ C for 3-5 days, as described (Cai et al., 2017).

### Antibody staining

Embryos were heat-fixed for immunostaining to ensure a low background during staining. Rabbit-anti-Ftz-antibodies were a gift from Markus Noll, rabbit-anti-Caudal-antibodies from Paul MacDonald, rabbit-anti-Stau from Daniel St. Johnston and mouse-anti-Osk from Zhenping Chen. Rat-anti-Hunchback and goat-anti-Cad antibodies were from the Asian Distribution Center for Segmentation Antibodies (Mishima, Japan). The monoclonal antibody 2B8 against Eve was from DSHB. DAPI for nuclear staining was used at 0.5mg/ml.

### Data analysis

All images were recorded using Zeiss LSM 700 and 710 inverted confocal microscopes.

---

**Figure S1.**
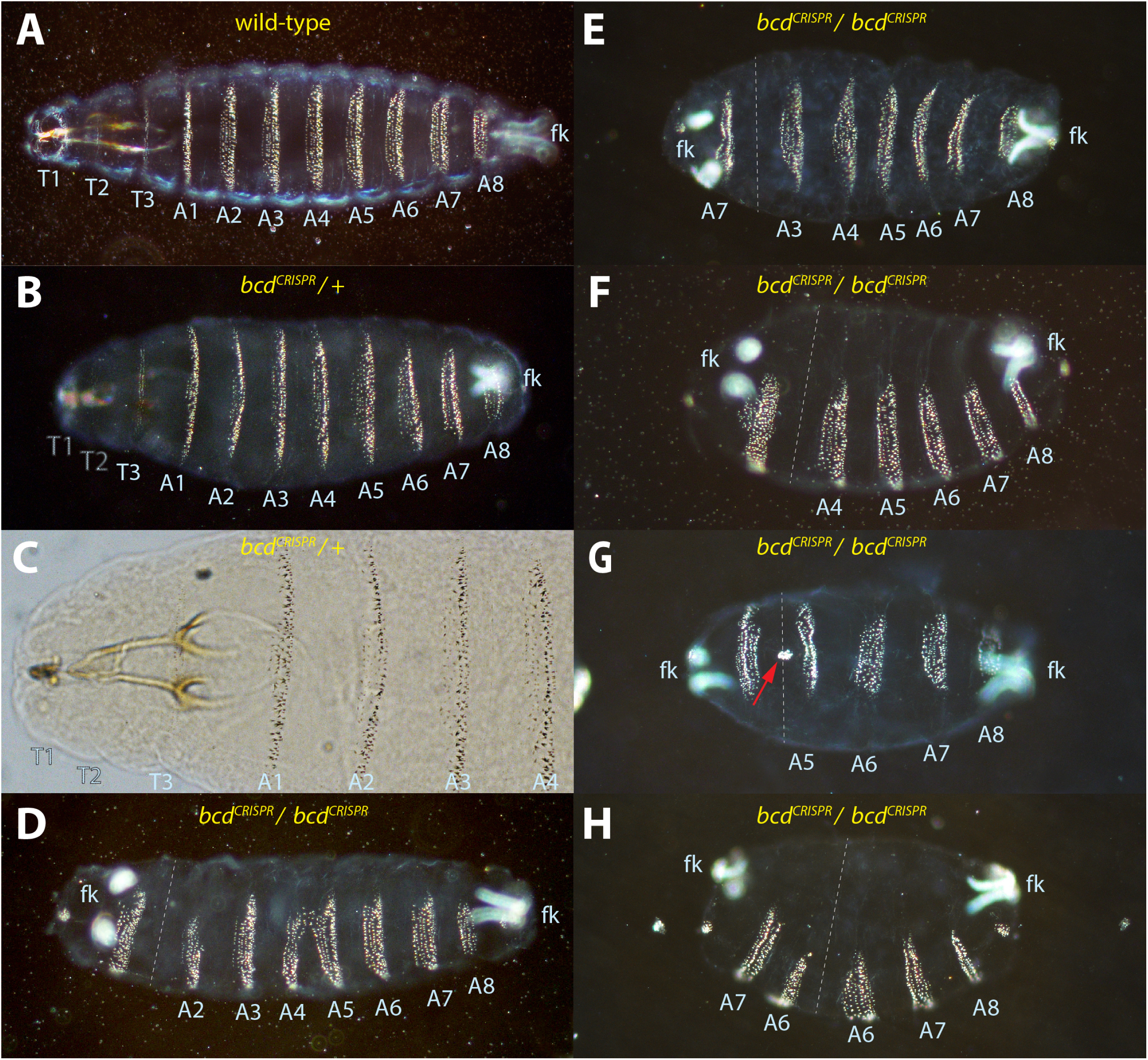
Phenotypic appearance of *bcd^CRISPR^* mutants. (A) cuticle of a wild-type larva as reference. (B) cuticle of a larva from heterozygous *bcd^CRISPR^* mothers and (C) anterior tip of the embryo in (B) to highlight the weak head phenotype with lack of cuticular patterning of T1 and T2. (D-H) cuticles from homozygous *bcd^CRISPR^* mothers, illustrating the phenotypic variability, with (G) representing the highest percentage for the phenotypic appearance. (D) cuticle where A2-A8 is present with a small posterior end of unknown segmental identity and the filzkörper (fk) duplicated to the anterior. (E) cuticle where A3-A8 is present with a small duplicated posterior end, A7 and the filzkörper. (F) cuticle where A4-A8 is present with a duplicated posterior end with unknown segmental identity and the filzkörper. (G) cuticle where A5-A8 is present with a duplicated posterior end, A7 and the filzkörper. (H) cuticle where A6-A8 is present with a duplicated posterior end, A7 and A6, and the filzkörper. Red arrow points towards a small circle of denticles infrequently observed, as also shown in Fig. 2L. All cuticles are oriented anterior to the left and showing the ventral side, unless otherwise noted. Identities of segments are indicated as abbreviations wherever identification was possible.

